# Profiling Siglec-7 and Siglec-9 ligands across the LuCaP PDX series: Implications for glyco-immune checkpoint inhibition in advanced prostate cancer

**DOI:** 10.64898/2026.09.02.748828

**Authors:** Ziqian Peng, Libby Blencoe, Kirsty Hodgson, Margarita Orozco-Moreno, Lizhi Cao, Wayne Gatlin, Li Peng, James Broderick, Richard Beatson, Jennifer Munkley

**Author notes:** Correspondence to: Jennifer Munkley.

## Abstract

Immunotherapies have shown limited efficacy in prostate cancer, largely due to its immunosuppressive tumour microenvironment. Hypersialylation contributes to immune evasion by engaging sialic acid-binding immunoglobulin-like lectins (Siglecs) on immune cells, forming glyco-immune checkpoints. Although this pathway represents a promising therapeutic target, the distribution of Siglec ligands across diverse prostate cancer phenotypes remains poorly understood. Here, we utilised high-affinity engineered sialoglycan-binding reagents (HYDRA) to perform immunohistochemical profiling of Siglec-7/9 ligands across a panel of 40 LuCaP patient-derived xenograft (PDX) models. Ligand expression was evaluated in relation to AR status and neuroendocrine phenotype. To determine the impact of androgen deprivation on the tumour glycome, ligand expression was compared between matched PDX lines grown in intact and castrated mice. Our findings reveal widespread but heterogeneous expression of Siglec-7/9 ligands across the LuCaP cohort. Expression levels were comparable between AR-positive adenocarcinoma models and AR-negative neuroendocrine variants, demonstrating that this glyco-immune checkpoint is maintained across distinct prostate cancer lineages. Under castration conditions, glycan remodelling occurred in a model-dependent manner. To evaluate these therapeutic dynamics in human disease, individual trajectory analysis of paired clinical specimens from the Movember GAP1-UTMA cohort was performed, revealing patient-specific responses that phenocopy the model-dependent glycan remodelling observed across the LuCaP lines. These findings support the sialoglycan-Siglec axis as a lineage-independent target and suggest that strategies aimed at disrupting Siglec-mediated immune suppression may be most effective when combined with androgen deprivation therapy to enhance anti-tumour immunity.

## Introduction

Prostate cancer remains a leading cause of cancer related mortality globally, largely driven by the development of resistance to standard-of-care therapies (1, 2). While androgen deprivation therapy (ADT) is initially effective, advanced tumours frequently adapt and progress to castration resistant prostate cancer (CRPC) (1, 3). CRPC is characterised by profound cellular and molecular heterogeneity, including lineage plasticity and neuroendocrine differentiation, and has limited therapeutic options (4, 5). Compounding this challenge, the prostate cancer tumour microenvironment is notoriously immunosuppressive, severely limiting the efficacy of conventional immune checkpoint inhibitors like anti-PD-1 or anti-CTLA-4 therapies (6, 7). Consequently, there is an urgent need to uncover novel immune evasion pathways that can be targeted in advanced prostate cancer.

Alterations in tumour glycosylation have emerged as crucial drivers of immune evasion and disease progression (8–11). In particular, hypersialylation, the overexpression of sialic acid-containing carbohydrates (sialoglycans) on the cell surface, acts as a glyco-immune checkpoint (12–14). These cancer-associated sialoglycans interact with sialic acid-binding Ig-like lectins (Siglecs), a family of inhibitory receptors predominantly expressed on innate and adaptive immune cells, including natural killer (NK) cells, macrophages, and T cells (14, 15). The interaction between sialoglycans on tumour cells and Siglec receptors on immune cells activates inhibitory signalling pathways inside immune cells. These pathways involve immunoreceptor tyrosine-based inhibitory motifs (ITIMs), which suppress anti-tumour immune responses facilitating tumour growth and metastasis (9, 11, 15).

We recently identified the Siglec-sialoglycan axis as a critical and actionable vulnerability in advanced prostate cancer (16–20). Utilising high-affinity, engineered Siglec-based sialoglycan binding reagents (termed HYDRAs), we demonstrated that sialoglycan ligands engaging Siglec-7, and Siglec-9 are upregulated in primary and bone metastatic human prostate cancer tissues (19). High expression of these ligands directly correlates with bone metastasis and poor patient prognosis. Furthermore, we established a therapeutic proof-of-concept by showing that systemic treatment with an engineered human sialidase (E-612), which strips cell surface sialic acids, can suppress tumour growth, enhance immune cell infiltration, and prolong survival times in mice with metastatic prostate cancer (19). While our initial findings underscore the therapeutic potential of targeting the Siglec-sialoglycan axis in prostate cancer, the precise distribution of these checkpoints across the landscape of advanced prostate cancer has not yet been fully mapped, and it is unknown how standard systemic treatments, such as hormone deprivation, alter the presentation of these immunomodulatory glycans.

Because prostate tumours frequently respond to therapeutic stress by undergoing phenotypic reprogramming (21, 22), understanding the stability and plasticity of the sialoglycan-Siglec axis under hormone therapy is critical for determining how desialylation-based therapies can be most effectively integrated into current treatments. To address this knowledge gap, we performed a comprehensive landscape profiling of Siglec-7 and Siglec-9 ligands (using HYDRA-7 and HYDRA-9 reagents, provided by Palleon Pharmaceuticals (23)) across the robust, clinically annotated Washington Carcinoma of the Prostate (LuCaP) patient-derived xenograft (PDX) series (24, 25) Comprising 40 distinct models, the LuCaP panel serves as a benchmark for replicating the phenotypic diversity of human prostate cancer spanning AR-positive (AR+) adenocarcinomas, aggressive AR-negative (AR-) neuroendocrine prostate cancers (NEPC), double-negative variants, and identical PDX lines grown in intact versus castrated mouse models. To ensure these model-dependent dynamics reflect true evolutionary pathways in human disease, we matched these findings against paired clinical specimens from the Movember Global Action Plan 1 Unique Tissue Microarray (GAP1-UTMA) cohort via individual trajectory tracking. Our findings reveal that Siglec-7 and -9 ligands are expressed across diverse prostate tumour lineages irrespective of AR status and reveal that castration induces model-dependent remodelling of sialylated glycans. Furthermore, tracking of individual clinical specimens demonstrates that hormone therapy induces patient-specific sialoglycan alterations in human disease that mirror the divergent responses observed across the LuCaP lines. Together, these insights provide a rationale for combining targeted desialylation therapies with standard hormonal interventions to develop new therapies for advanced prostate cancer.

## Methods

### Immunohistochemistry

The HYDRA immunohistochemistry protocol has been previously published by us (19). HYDRA-7 and HYDRA-9 reagents, which correspond to ligand binding domains of their corresponding Siglecs (Siglec-7 and Siglec-9 respectively) (23), were provided by Palleon Pharmaceuticals (Waltham, MA). HYDRA reagents were diluted in Antibody Diluent (Dako, S302283-2). All slides were stained using the IHC Prep & Detect Kit for Rabbit/Mouse Primary Antibody (Proteintech, PK10019), following the manufacturer’s protocols. A Sodium Citrate Antigen Retrieval Buffer (Proteintech, PR30001) was used, and tissues were incubated with HYDRA-7 at 1.5 μg/mL or HYDRA-9 at 0.25 μg/mL for 1 hour before the secondary antibody incubation. HistoClear (National Diagnostics, HS-200) and HistoMount (National Diagnostics, HS-103) were used instead of Xylene and the mounting media from the kit. Images were acquired and processed using the ZEISS Axioscan 7 and ZEN Microscopy Software (blue edition) and analysed using QuPath (version 0.6) software as described by us previously (20, 26). Each PDX sample was analysed in triplicate from the same mouse and across three independent mice. Positive cells were outlined and classified as having low, moderate or high staining intensity to calculate a 0-300 HistoScore, calculated as the sum of each staining intensity score (1+, 2+, 3+) multiplied by the percentage of cells classified at that intensity using the equation staining Index (H-score) = Σ (each intensity score × % of cells at that intensity). In line with previous studies (19, 26–28), only epithelial cells were scored. A Histoscore <50 was considered negative, scores of 50-100 were weakly positive/negligible, scores >100 were recorded as positive, and scores of >200 were recoded as highly positive.

### Tissue Microarrays and Clinical Cohort

The LuCaP PDX TMA contains 40 previously published PDX model tumours (24, 25), established from distinct patients from specimens, acquired at either radical prostatectomy or at autopsy. Tissues were implanted and grown subcutaneously in intact or castrated immunocompromised mice (in triplicate). A summary of the specific LuCaP models utilised in this study, including their clinical and phenotypic characteristics, is provided in Table 1. Castration resistant derivatives of each LuCaP model were created by enabling the tumours to grow and progress in castrated mice. Strict core selection protocols were applied to ensure data quality; areas of necrosis, apoptosis, or mouse stromal contamination were actively avoided during core punching, and any remaining areas of necrosis were strictly excluded from the final immunohistochemical analysis. Cores were 5 µm thick, 1.5 mm diameter, and were included in triplicate per LuCaP line. To monitor Siglec-7 ligands in human disease under therapeutic stress, clinical samples from the Movember Global Action Plan 1 Unique Tissue Microarray (GAP1-UTMA) cohort were evaluated (29); HYDRA-7 immunohistochemistry data for this cohort has been previously analysed and published by us (19).

### Statistical analysis

Statistical analyses were conducted using the GraphPad Prism software (version 11) using suitable tests as described in each figure legend. Data are presented as the mean of three independent samples ± standard error of the mean (SEM). Statistical significance is denoted as *p<0.05, **p<0.01, ***p<0.001 and ****p<0.0001.

## Results

### 1. Comprehensive profiling reveals heterogeneous Siglec-7 and Siglec-9 ligand expression across LuCaP PDX models

To define the landscape of Siglec ligand expression in advanced prostate cancer, we performed immunohistochemical profiling of Siglec-7 and Siglec-9 ligands across a panel of 40 LuCaP PDX models encompassing the molecular and phenotypic diversity of advanced disease (24, 25) (**Table 1**). Siglec ligands were detected using the novel HYDRA reagents (provided by Palleon Pharmaceuticals), including HYDRA-7 and HYDRA-9, which comprise multimeric Siglec domains with enhanced avidity for the detection of sialoglycans recognised by Siglec-7 and Siglec-9, respectively (23, 30). These reagents have previously been validated by us for use on prostate cancer tissues (19). Siglec-7 and Siglec-9 ligands were detected across the majority of LuCaP models, although the intensity of staining varied considerably between tumours (**Figure 1A**). While some models, including LuCaP 70 and LuCaP 136, exhibited strong epithelial staining for both ligand types, other models, including LuCaP 96 and LuCaP 189.3, displayed low levels or absent ligand expression (**Figure 1B**). Notably, the abundance of Siglec-7 ligands did not consistently correlate with Siglec-9 ligands within individual models, highlighting distinct, patient-specific patterns of tumour sialylation. For instance, LuCaP 96CR and LuCaP 145.2 demonstrated high Siglec-7 binding but low Siglec-9 levels (**Figure 1C**), whereas LuCaP 23.1 and LuCaP 35 exhibited the opposite profile (**Figure 1D**). Although ligand abundance differed considerably between individual PDX models, both Siglec-7 and Siglec-9 ligands were consistently represented across the panel, suggesting that hypersialylation and the associated Siglec glyco-immune checkpoint are common features of advanced prostate cancer. Together, these findings provide a comprehensive atlas of Siglec-7 and Siglec-9 ligand expression across the LuCaP PDX panel and demonstrate that the Siglec glyco-immune checkpoint is broadly, but heterogeneously, represented across advanced prostate cancer.

**Figure 1.**
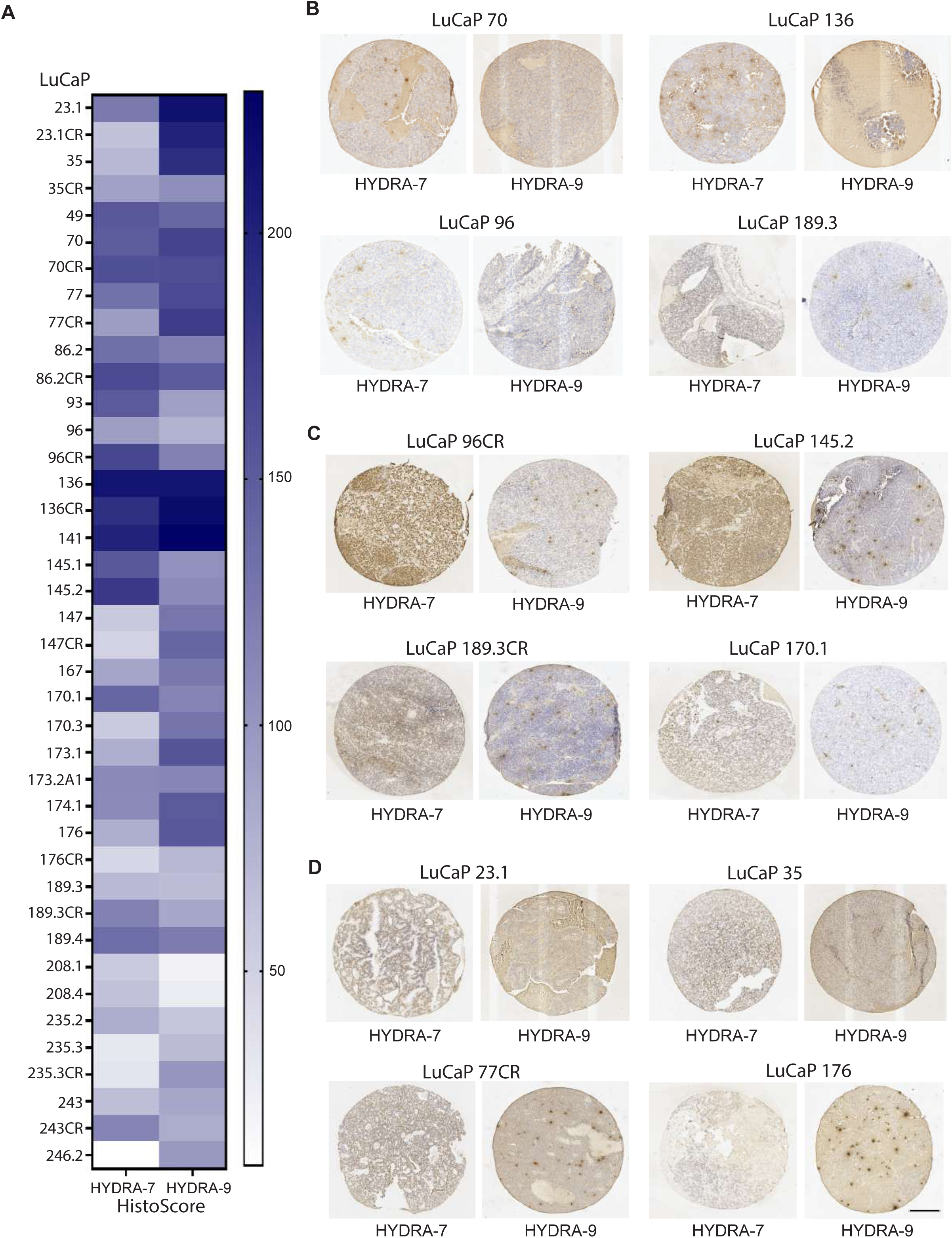
Expression of Siglec-7 and Siglec-9 ligands across the LuCaP PDX models. (**A**) Heatmap illustrating the Histoscores for Siglec-7 and Siglec-9 sialoglycan ligands (monitored using HYDRA-7 and HYDRA-9 immunohistochemistry). (**B**) HYDRA immunohistochemistry analysis shows both Siglec-7 and Siglec-9 ligands are expressed at high levels in LuCaP 70 and LuCaP 136, but at low levels in LuCaP 96 and LuCaP 189.3. (**C**) For LuCaP 96CR, 145.2, 189.3CR, and 170.1 Siglec-7 ligands are expressed at relatively higher levels than Siglec-9 ligands. (**D**) For LuCaP 23.1, 35, 77CR and 176 Siglec-9 ligands are expressed at relatively higher levels than Siglec-7 ligands. Scale bar is 200 µm.

### 2. Siglec-7 and Siglec-9 ligands are maintained across AR-positive and AR-negative prostate cancer

To investigate whether Siglec ligand expression correlates with AR status, LuCaP PDX models were stratified into AR-positive and AR-negative groups, and Siglec-7 and Siglec-9 ligand expression was compared. As AR loss is frequently associated with lineage plasticity and neuroendocrine differentiation, this analysis sought to establish whether the Siglec glyco-immune checkpoint is preserved across molecular phenotypes of advanced prostate cancer. Siglec-7 and Siglec-9 ligands were detected in both AR-positive and AR-negative PDX models, with tumours from each group showing comparable staining intensity, suggesting that ligand expression is retained despite the loss of AR expression (**Figure 2A,B**). Interestingly, while Siglec-7 ligand levels remained highly consistent between the groups, a noticeable trend towards reduced Siglec-9 ligand expression was observed in the AR-negative models; however, this decrease did not reach statistical significance. Although inter-model heterogeneity was evident within both groups, no obvious differences in staining patterns were observed between AR-positive and AR-negative tumours. These findings suggest that AR status does not definitively influence the abundance of Siglec-7 or Siglec-9 ligands within advanced prostate cancer. The maintenance of Siglec-7 and Siglec-9 ligands across both AR-positive and AR-negative disease states suggests that the Siglec glyco-immune checkpoint is conserved despite profound changes in tumour phenotype, supporting its potential as a therapeutic target across the spectrum of advanced prostate cancer.

**Figure 2.**
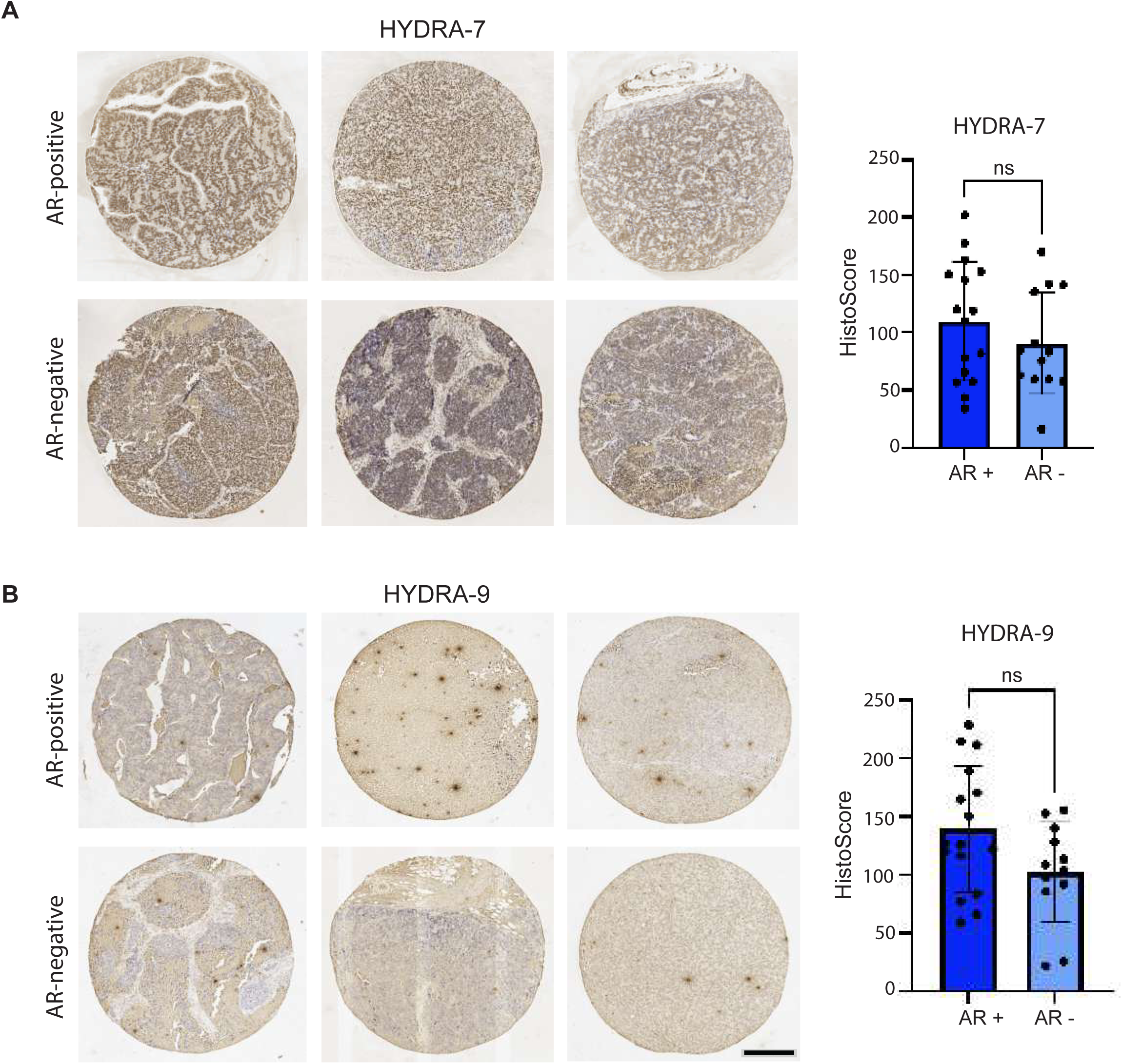
Advanced prostate cancer maintains Siglec-7 and Siglec-9 ligand expression across both AR-positive (AR+) and AR-negative (AR-) lineages. (**A**) HYDRA-7 immunohistochemistry analysis shows Siglec-7 ligands are expressed at similar levels in both AR+ prostate adenocarcinomas and AR- prostate tumours (n=29, unpaired t test, p=0.3006). (**B**) HYDRA-9 immunohistochemistry shows Siglec-9 ligands are expressed in both AR+ prostate adenocarcinomas and AR- NEPC and double negative prostate tumours, with a noticeable but not statistically significant reduction of Siglec-9 ligands in AR negative lines (n=29, unpaired t test, p=0.0618). Representative images are shown. AR- tumours include both NEPC and double negative tumours. Scale bar is 200 µm.

### 3. Androgen deprivation remodels the Siglec glyco-immune checkpoint in a heterogeneous, patient-specific manner

Given the central role of androgen signalling in prostate cancer biology, we next investigated whether androgen deprivation alters tumour-associated Siglec ligand expression. Matched LuCaP PDX models were evaluated following growth in intact and castrated mice, enabling assessment of glycome changes induced by androgen withdrawal within the same tumour context. Growth in castrated conditions resulted in model dependent changes in both Siglec-7 and Siglec-9 ligand expression, suggesting that androgen withdrawal drives tumour-specific remodelling (**Figure 3 and 4**). We identified three distinct patterns of response across the cohort. Several models, such as LuCaP 23.1 for Siglec-7 and LuCaP 35 for Siglec-9, demonstrated reduced ligand expression in androgen depleted conditions. This pattern is consistent with partial androgen dependence of tumour sialylation pathways. In contrast, other models showed increased ligand expression when grown in castrated mice. Notably, LuCaP 96 exhibited upregulation of both Siglec-7 and Siglec-9 ligands under castrated conditions. These increases suggest that therapeutic stress can promote adaptive remodelling of the tumour glycome, potentially promoting inhibitory sialoglycan-Siglec immune interactions. A third group of models, including LuCaP 136 (Siglec-7) and LuCaP 23 (Siglec-9), displayed minimal alteration in ligand expression. This indicates that Siglec ligand expression can also be maintained independently of androgen signalling.

**Figure 3.**
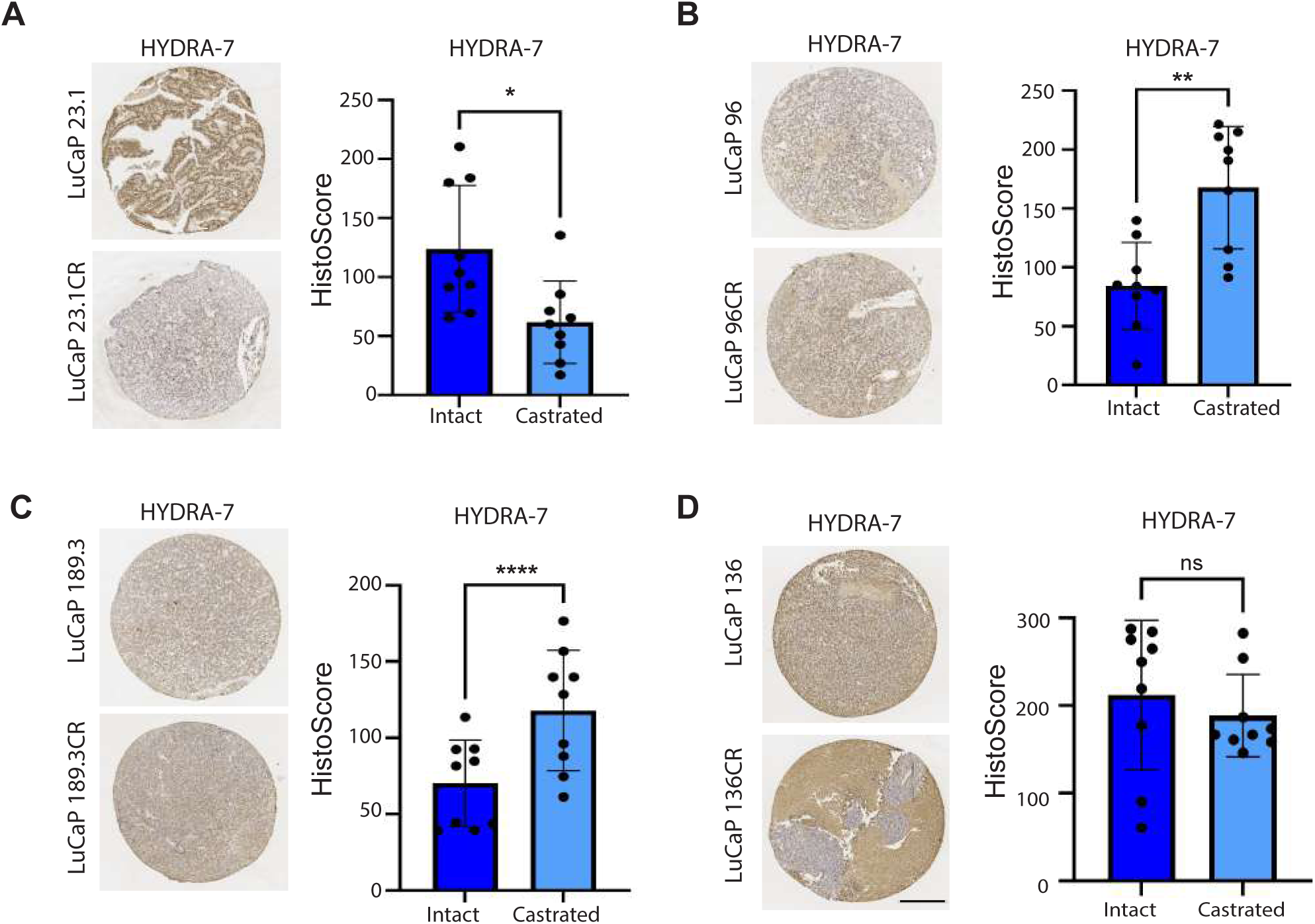
Androgen deprivation induces heterogeneous, patient-specific remodelling of Siglec-7 ligand expression in LuCaP PDX models. (**A**) LuCaP 23.1 exhibits a significant reduction in Siglec-7 ligand expression in PDX samples grown in castrated versus intact mice (unpaired t-test, p=0.0103). (**B**) In contrast, LuCaP 96 displays a significant increase in Siglec-7 ligand abundance in castrated mice compared to intact controls (unpaired t-test, p=0.0012). (**C**) LuCaP 189.3 similarly demonstrates a significant increase in Siglec-7 ligand expression following host castration (unpaired t-test, p=0.0001). (**D**) LuCaP 136 shows no significant change in Siglec-7 ligand levels between intact and castrated conditions (unpaired t-test, p=0.4793). Representative HYDRA-7 immunohistochemistry images are shown for each model (each PDX sample was analysed in triplicate across 3 mouse models). Scale bar = 200 µm.

**Figure 4.**
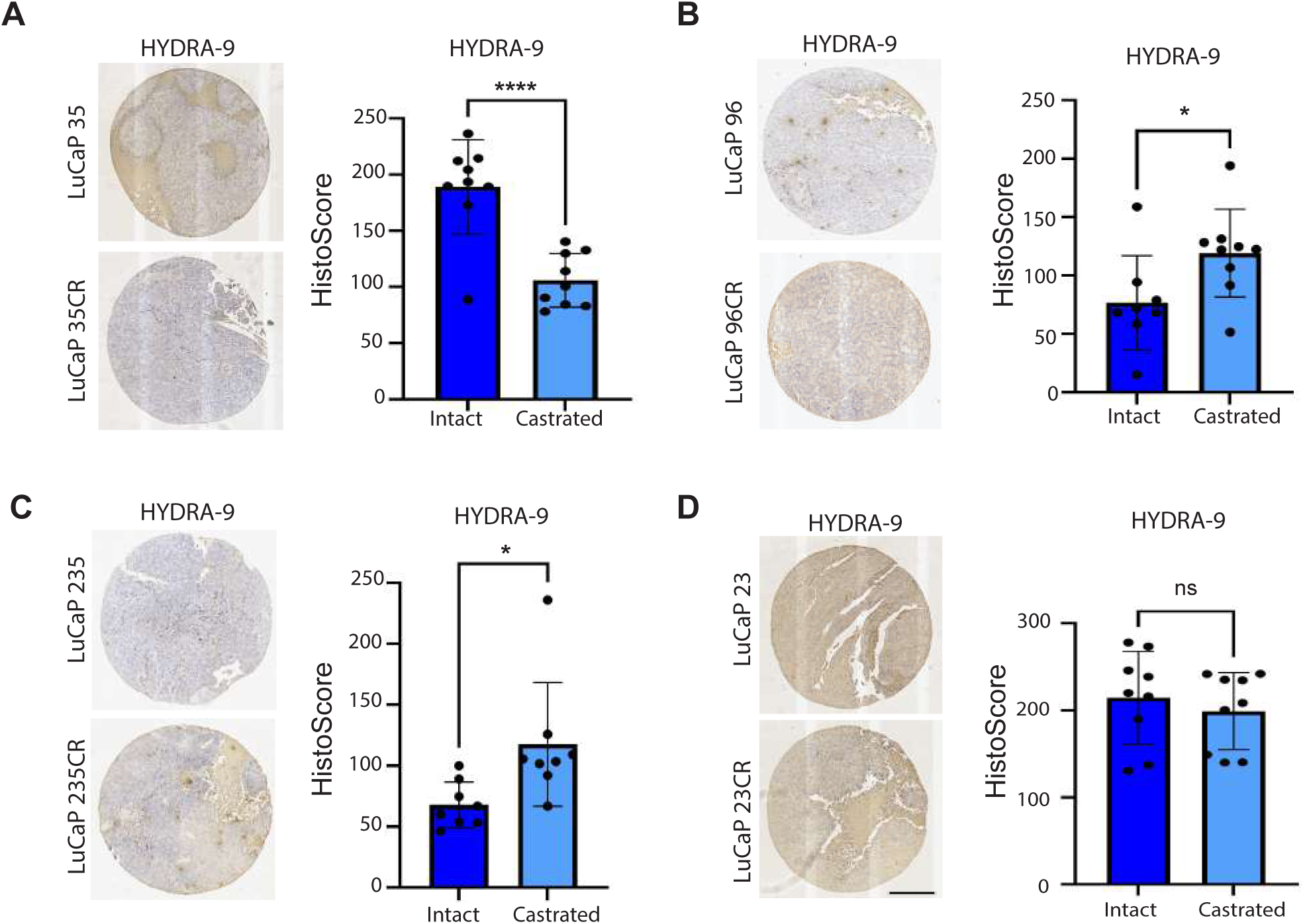
Androgen deprivation induces heterogeneous, patient-specific remodelling of Siglec-9 ligand expression in LuCaP PDX models. (**A**) LuCaP 35 shows a significant reduction in Siglec-9 ligand expression in PDX samples grown in castrated versus intact mice (unpaired t test, p=0.0002). (**B**) In contrast, LuCaP 96 displays a significant increase in Siglec-9 ligands in castrated mice compared to intact mice (unpaired t test, p=0.0405). (**C**) LuCaP 235 similarly demonstrates a significant increase in Siglec-9 ligand expression in androgen depleted conditions (unpaired t test, p=0.0211). (**D**) LuCaP 23 shows no significant change in Siglec-9 ligand levels between samples grown in intact and castrated conditions (unpaired t test, p=0.5161). Representative HYDRA-9 immunohistochemistry images are shown for each model (each PDX sample was analysed in triplicate across 3 mouse models). Scale bar = 200 µm.

To investigate whether these model-dependent dynamics reflect evolutionary dynamics in human disease, we re-evaluated our previously published clinical profiling data for Siglec-7 ligands in the Movember Global Action Plan 1 Unique Tissue Microarray (GAP1-UTMA) cohort (29). While our previous analysis showed no statistically significant overall change when comparing Siglec-7 ligands in paired pre-ADT and CRPC tissues, we hypothesised that by looking at the overall cohort we may have missed patient-specific changes. Re-evaluation of our published data (19) to track individual patients within the paired dataset, revealed a context-dependent glycan profile which replicates the model-dependent heterogeneity observed in our LuCaP PDX castration studies (**Figure 5**). Upon progression to castration resistance following clinical ADT, a subset of patients showed upregulation of Siglec-7 ligands, a second subset exhibited downregulation, and a third group showed no change in ligand abundance. As the Movember GAP1-UTMA project is no longer active, it was not possible for us to access additional GAP1-UTMA slides, meaning we have been unable to monitor ligands for Siglec-9 in these patient cohorts. Together, these findings establish that androgen deprivation dynamically remodels the Siglec glyco-immune checkpoint in a patient-specific manner. Although Siglec-7 and Siglec-9 ligands are broadly maintained across prostate cancer phenotypes, therapeutic pressure can induce divergent alterations in ligand expression. This variability highlights the importance of understanding treatment-induced glycan changes when developing strategies targeting the sialoglycan-Siglec axis. Furthermore, it supports biomarker guided combination approaches incorporating desialylation-based therapies with androgen deprivation.

**Figure 5.**
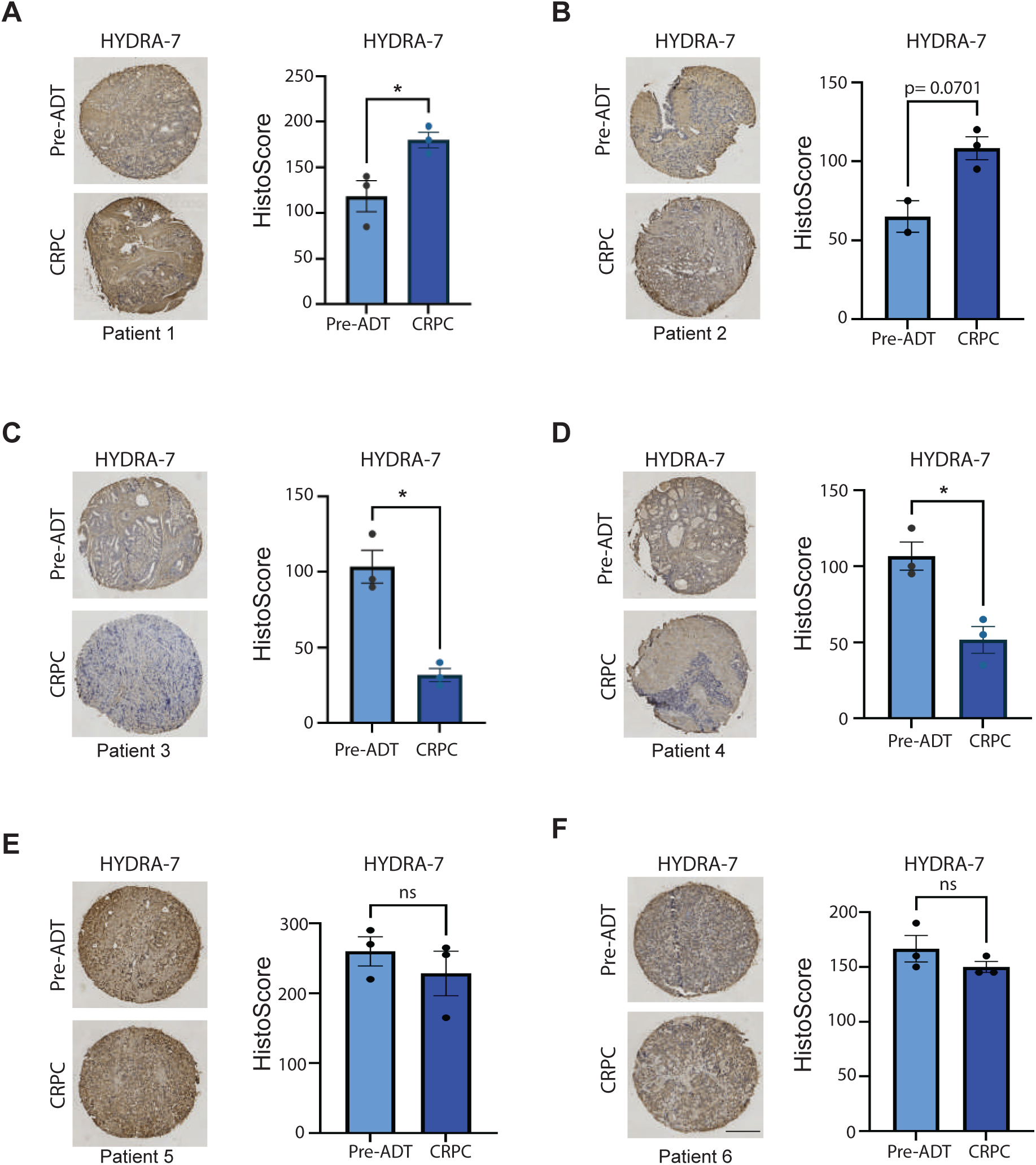
Individual trajectory tracking in human clinical specimens suggests patient-specific sialoglycan remodelling in CRPC. HYDRA-7 immunohistochemistry analysis of Siflec-7 ligands in matched pre-ADT and CRPC samples from the Movember Global Action Plan 1 Unique Tissue Microarray (GAP1-UTMA) cohort (29). (**A**) For patient 1 there is a significant increase in Siglec-7 ligands in CRPC samples compared to matched pre-ADT samples from the same patient (Welsh’s test, p=0.0481). (**B**) For patient 2, Siglec-7 ligands were also higher in matched CRPC tissues (Welsh’s test, p=0.0701). (**C**,**D**) For patient 3 and 4, Siglec-7 ligands were significantly lower in CRPC tissues compare to matched pre-ADT samples (Welsh’s t tests, p=0.0129 and p=0.0128). (**E**,**F**) For patients 5 and 6, there was no overall change in Siglec-7 levels following progression to CRPC (Welsh’s t tests, p=0.4585 and p=0.3003). Representative images are shown for each patient. Scale bar is 200 µm.

## Discussion

Advanced prostate cancer is defined by substantial molecular and cellular heterogeneity, which enables rapid tumour adaptation to therapeutic pressure through mechanisms such as AR pathway rewiring, lineage plasticity, and neuroendocrine differentiation (1, 31). Although hormone therapy remains the cornerstone of treatment for advanced disease, progression to CRPC is frequently accompanied by loss of dependence on canonical AR signalling and emergence of highly aggressive variant tumour states (32). Identifying therapeutic vulnerabilities that remain during this evolutionary trajectory is an urgent unmet need (33). In this study, we demonstrate that the Siglec-7/9 glyco-immune checkpoint is broadly maintained across a comprehensive panel of advanced prostate cancer phenotypes, spanning AR-positive adenocarcinomas and AR-negative NEPC variants. Furthermore, we show that androgen deprivation induces heterogeneous, patient-specific remodelling of Siglec ligand expression, revealing dynamic regulation of the tumour glycome during therapeutic adaptation.

Aberrant glycosylation is increasingly recognised as a fundamental hallmark of cancer biology, influencing tumour proliferation, invasion, metastasis, immune evasion, and response to therapy (10, 34, 35). Prostate cancer displays extensive alterations in glycan processing, with changes in glycosyltransferase expression and sialylation patterns contributing to disease progression (16, 36, 37). Importantly, androgen signalling has previously been implicated in regulating tumour-associated glycosylation, with AR activity shown to modulate glycosylation related pathways, indicating that the prostate cancer glycome can be dynamically influenced by hormonal signalling (36, 38–40). Prior studies established a functional link between androgen signalling, ST3Gal1-mediated sialoglycan synthesis, and the generation of Siglec-7/9 ligands that suppress anti-tumour immune responses (41). Our current findings extend these observations by demonstrating that androgen deprivation does not produce a uniform glycomic response across advanced prostate cancer models. Instead, androgen withdrawal results in divergent patterns of Siglec ligand remodelling, reflecting the complex regulation of glycan biology in treatment adapted disease that extends beyond classical AR control.

A key finding of this study is that Siglec-7 and -9 ligand expression is maintained across prostate cancer lineage states. While lineage plasticity and neuroendocrine differentiation represent major escape routes from AR-targeted therapies, these transitions can result in the loss of conventional therapeutic targets, including the AR itself (32). Our finding that AR-positive adenocarcinomas and AR-negative NEPC and double negative models retain comparable levels of Siglec-7 and Siglec-9 ligands suggests that this glyco-immune checkpoint is a lineage-independent vulnerability in advanced prostate cancer. Consequently, unlike targets that are restricted to a particular molecular subtype, the sialoglycan-Siglec axis may remain therapeutically relevant even following escape from androgen dependence, providing a broad window for therapeutic intervention across heterogeneous disease states.

The relationship between androgen signalling and Siglec ligand expression is clearly more complex than a linear AR-dependent regulatory pathway. Following host castration, we identified three distinct patterns of glycomic adaptation across the LuCaP series: increased Siglec ligand abundance, reduced Siglec ligand expression, and relative stability. Notably, we also observed substantial variation in ligand staining intensity between individual cores within the same LuCaP models. This intra-model variability likely reflects a degree of intra-tumoural spatial heterogeneity, suggesting that Siglec ligand expression is likely not uniform across the geographic landscape of a single tumour. Models exhibiting upregulation of Siglec ligand abundance following androgen deprivation may exploit an adaptive, stress-induced program to promote immune evasion. Conversely, models showing reduced ligand expression suggest that a subset of advanced tumours may retain dependence on androgen-regulated glycosylation machinery. Finally, the cohort displaying stable ligand expression indicates that Siglec ligand presentation can become completely uncoupled from androgen status. Importantly, because these three divergent patterns occurred across both AR-positive and AR-negative phenotypes, baseline AR status alone is insufficient to predict how a patient’s glycome will respond to androgen withdrawal. When we applied individual trajectory tracking to the paired human clinical specimens from the Movember GAP1-UTMA cohort, we detected divergence in sialoglycan expression following progression to CRPC. This approach also identified clinical subpopulations with Siglec-7 ligand upregulation, downregulation, or stable expression, suggesting that the heterogeneous expression patterns revealed by our PDX lines are also observed in human disease.These findings have important implications for understanding treatment adaptation in prostate cancer. ADT is known to remodel tumour cell biology and the tumour microenvironment (42, 43); however, its impact on immune-regulatory glycans remains incompletely understood. Our data suggest that androgen deprivation may influence immune vulnerability through the modulation of tumour-associated sialoglycans. In some tumours, ADT may reduce Siglec ligand-mediated immune suppression, whereas in others it may promote an adaptive increase in inhibitory glycan expression. This heterogeneity may contribute to variable responses observed when combining hormonal therapies with immune-directed approaches and highlights the need to consider treatment-induced glycomic changes when designing combination strategies.

Exploiting tumour glycosylation represents a powerful therapeutic strategy, supported by our previous demonstration of the anti-tumour efficacy of E-612 in prostate cancer models, which established that targeting Siglec-engaging sialoglycans can suppress prostate cancer progression and bone metastasis by overcoming the protective, sialic acid-dependent immune barrier (19). This therapeutic approach aligns with previous studies demonstrating that hypersialylation drives profound immune escape and that removing surface sialic acids can repolarise the immunosuppressive tumour microenvironment to boost anti-tumour immunity (44–46). Notably, Palleon Pharmaceuticals has progressed their B7-H3 targeted sialidase fusion protein (47) into clinical trials for platinum-resistant ovarian cancer (NCT07541534). As the immune checkpoint protein B7H3 is highly expressed on the surface of the most lethal prostate tumours (48), we propose that this therapeutic strategy could be applied to leverage enzymatic sialoglycan degradation to specifically target advanced prostate cancer cells. In line with this approach, the current study shows that Siglec-7 and -9 ligands are maintained in advanced prostate cancer despite lineage transitions and treatment resistance. The continuous presence of these actionable targets across the LuCaP cohort underscores that glycan-targeted therapies are likely a viable therapeutic option across the full spectrum of advanced prostate cancer evolution.

While the LuCaP PDX series provides insight into the phenotypic and lineage diversity of advanced clinical prostate cancer (summarised in Table 1), certain limitations of this model system must be acknowledged, including the use of immunocompromised murine models, which lack a functional human immune system. Furthermore, while HYDRA profiling shows the expression levels of Siglec-7/9 ligands, this pathway relies on engagement with inhibitory receptors on immune cells, which have not been investigated in this study. Finally, while the LuCaP series originates from diverse clinical metastatic sites (including bone and visceral tissue), growing these models subcutaneously in mice may not fully replicate the complex microenvironmental factors, such as the unique tumour microenvironment and cytokine profile of human metastases, which likely influence global tumour sialylation patterns. To address this, future studies utilising humanised mouse models and/or ex vivo patient-derived explant cultures will be performed to fully characterise how alterations to sialoglycans may influence the response to targeted sialidase therapies.

In conclusion, this study identifies the Siglec-7/9 glyco-immune checkpoint as a conserved feature of advanced prostate cancer that persists across AR-dependent and AR-independent disease states. We demonstrate that androgen deprivation dynamically remodels this checkpoint in a tumour-specific manner, revealing glycomic plasticity as an additional mechanism of therapeutic adaptation. Future studies integrating glycomic and spatial immune cell profiling will be essential to map the precise glycoprotein carriers driving this plasticity, and to define how treatment induced glycan remodelling alters the clinical prostate cancer tumour immune microenvironment. Together with our previous findings targeting tumour sialylation in prostate cancer, these data support development of combination strategies that exploit glycan vulnerabilities alongside androgen-directed therapies to improve treatment responses in advanced disease.

## Conflict of Interest

LC, WG, LP and JB are employees and hold stock in Palleon Pharmaceuticals.

## Acknowledgements

We thank the patients and their families, Pete Nelson, Evan Yu, Heather Cheng, Bruce Montgomery, Jessica Hawley, Mike Schweizer, Daniel Lin, Funda Vakar-Lopez, Michael Haffner, Martine Roudier, Lawrence True, Meagan Chambers, Colm Morrissey, and the rapid autopsy teams for their contributions to the University of Washington Medical Center Prostate Cancer Donor Rapid Autopsy Program. The authors would also like to thank Professor Eva Corey and Professor Colm Morrissey for kindly providing the LuCaP TMAs for analysis in our study.

## Author Contributions

Z.P performed immunohistochemistry analysis of clinical tissues. Z.P and L.B performed data analysis. L.C., W.G., L.P and JB. provided the HYDRA reagents for the study. Z.P., J.M. and K.H. verified the underlying data. J.M., K.H., and Z.P. jointly designed, analysed, and interpreted the study. J.M. and Z.P. wrote the original manuscript draft and made the figures. L.C, J,B,, W.G., L.P, K.H and R.B. contributed to the critical review of the manuscript. J.M. contributed to funding acquisition. J.M, K.H, MOM and R.B. contributed to project supervision. All authors read the manuscript, agreed with the content, and were given the opportunity to provide input.

## Ethics approval

All human studies were reviewed by the appropriate ethics committee and performed in accordance with the ethical standards laid down in the Declaration of Helsinki. All autopsy tissues were collected from patients who had signed written informed consent under the aegis of the Prostate Cancer Donor Program at the University of Washington (25). The IRB of the University of Washington approved this study. All PDX experiments were approved by the University of Washington IACUC. For animal experiments, the ‘Principles of Laboratory Animal Care’ were followed as well as specific national laws as detailed in previous publications (24, 25).

## Funding

This work was funded by Prostate Cancer UK [RIA21-ST2-006], the Medical Research Council [MR/R015902/1], the JGW Patterson Foundation, and Prostate Cancer Research (grant reference 6974). This work is also supported by the Pacific Northwest Prostate Cancer SPORE (P50CA97186), the PO1 NIH grant (PO1 CA163227), the Prostate Cancer Foundation, and the Institute for Prostate Cancer Research (IPCR).

## Data Availability Statement

The data that support the findings of this study are available from the corresponding author upon reasonable request.

## Notes

### Competing Interest Statement

LC, WG, JB and LP are employees and hold stock in Palleon Pharmaceuticals.

### Summary of Updates

Additional analysis of clinical specimens (Figure 5)

